# *In Vivo* Measurements of the Frequency-Dependent Impedance of the Spinal Cord

**DOI:** 10.1101/252965

**Authors:** Marcel Utz, John W. Miller, Chandan G. Reddy, Saul Wilson, Kingsley O. Abode-Iyamah, Douglas C. Fredericks, George T. Gillies, Matthew A. Howard

## Abstract

Improved knowledge of the electrode-tissue impedance will be useful in optimizing the clinical protocols and resulting efficacy of the existing and emerging approaches to spinal cord stimulation. Toward that end, the complex impedance (amplitude and phase) of *in vivo* ovine spinal cord tissue was measured at the electrode-pial subdural surface interface from 5 Hz to 1 MHz, and with the bi-polar electrodes oriented both parallel and perpendicular to the rostral-caudal axis of the spinal cord. At stimulation frequencies above 10 kHz, most of the impedance then becomes resistive in nature and the phase difference between the stimulation signal and the resulting current drops to ≈ 10˚, thus maximizing power transfer to the tissues. Also, at these higher frequencies, the current pulse maintains significantly greater fidelity to the shape of the stimulation signal applied across the electrodes. Lastly, there were lower impedances associated with parallel as opposed to perpendicular orientation of the electrodes, thus reflecting the effects of fiber orientation within the spinal cord. Impedance differences of this kind have not been reported with epidural stimulation because of the electrical shunting effects of the intervening layer of relatively high conductivity cerebrospinal fluid. These observations provide a quantitative basis for improved models of spinal cord stimulation and suggest certain advantages for direct intradural stimulation relative to the standard epidural approaches. (Some figures in this article are in colour only in the electronic version)

## 1. Introduction

One of the primary factors limiting the clinical efficacy of epidural spinal cord stimulation (SCS) is the presence of the relatively high electrical conductivity cerebrospinal fluid (CSF), *σ* ≈ 1.7 (Ω•m)^-1^, which fills the space between the underside of the dura and the pial surface of the spinal cord. Because the current densities produced by the stimulator’s electrodes are largely shunted through the CSF, only a thin surface layer of axons (approximately 250 μm deep from the dorsal pial surface of the spinal cord) can typically be activated [1]. Moreover, the therapeutic window is very small: increasing the strength of the stimulation signals in order to excite axons deeper within the spinal cord results in painful spillover excitation of non-targeted structures such as the dorsal nerve rootlets. In response to these limitations, many substantial efforts have been undertaken for purposes of modeling the stimulation process, optimizing the designs of the implanted leads, and synthesizing improved excitation montages, as discussed in the reviews of Bradley [2], North [3] and Levy [4]. See also the early papers of Holsheimer and colleagues, for example [5,6,7,8,9,10,11], who developed many of the computational tools needed for that work.

More recently, several investigators have begun exploring neurophysiological effects associated with high frequency (> 1 kHz) stimulation [12,13,14]. Initial clinical trials of epidural stimulation at frequencies up to 10 kHz have demonstrated patient preference for this mode of stimulation in the short term [15], along with improved long term outcomes [16,17,18] relative to the low frequency norms in which the therapy ultimately fails in up to half of all patients, e.g. [19]. Efforts are now underway to help elucidate the mechanisms of therapeutic action for high frequency stimulation effects [20], in concert with suggestions for further improvement in clinical strategies [21].

These important advances notwithstanding, a number of fundamental issues associated with epidural placement of the leads and the shunting effects of the CSF remain unresolved, including lead migration, stimulator power consumption, and selectivity and depth of coverage within the neural tissues. In fact, it was with these limitations in mind that we suggested the possibility of direct stimulation of the pial surface of the spinal cord [22,23,24], began developing the technologies needed to enable it [25,26], and carried out preliminary tests of this approach (termed the “I-Patch”) in a large animal model [27,28]. Part of the rationale for this intradural design is that when targeting spinal cord structures, direct stimulation using stimuli of any frequency should be able to achieve therapeutic neural activation at much lower power levels by circumventing the CSF shunting effects. Moreover, the question was open as to whether the stimulus pattern produced by a high frequency pulse train delivered epidurally would degrade in fidelity after passing through the intervening CSF as opposed to a signal applied directly to the spinal cord.

In order to investigate these points and then optimize the design of an intradural direct SCS system, knowledge of the frequency-dependent impedance at the interface between the electrode and the pial surface is needed. While measurements of that type have been made for deep brain stimulators [29], only limited types of data have been taken for spinal cord at or near the electrode-epidural tissue interface [30,31]. Therefore, we designed an experiment using our ovine model of intradural stimulation [28,29] in which we place a pair of electrodes directly on the pial surface of the spinal cord of adult sheep, with the intent here to record the applied voltage and resulting current waveforms as a function of frequency. This would then allow us to determine the frequency-dependent impedance and phase at that interface and extract the resistive and reactive components of impedance. By aligning the electrode pair along the rostral-caudal axis of the spinal cord and then perpendicular to it, we would also be able to investigate how axonal fiber orientation within the spinal cord affects those quantities. In what follows, we describe the details of the experimental protocol, present the results of the measurements, and then discuss their meaning and impact within the context of both epidural and intradural direct spinal cord stimulation.

## 2. Methods

This investigation was part of an overall study on the potential efficacy of SCS on ovine models of spinal cord injury and neuropathic pain, approved by the Institutional Animal Care and Use Committee (IACUC) of the University of Iowa (U.I.) under IACUC Approval No. 1308149. The intradural stimulation experiments described below were carried out on three of the adult sheep (≈ 70 kg weight) in that study, during the terminal surgical procedure for each animal. The methods for induction and maintenance of anesthesia for the surgeries, and then for euthanasia following the stimulation experiments, were as used in our previous trials of ovine intradural SCS and are described in detail elsewhere [27,32].

### 2.1 Surgical procedure

The neurosurgical steps consisted of exposure of the spinal column at the mid-thoracic level, followed by laminectomy and durotomy. After the thecal sac was opened, a micropositioner was used under neuroendoscopic guidance to position a pair of hemispherical electrodes onto the dorsal pial surface of the spinal cord. The electrodes were 1.0 mm in diameter, the center-to-center distance between them was 2.0 mm, and the depth of indentation into the spinal cord insertion was ≈ 0.2 mm. Stimulation measurements were made with the alignment of the electrodes along the rostral-caudal axis of the spinal cord (i.e., in the “parallel” position), and at 90˚ with respect to that axis (i.e., in the “perpendicular” position). A close-up photo of the electrodes on the distal tip of the probe is shown in figure 1(a) and its location on the exposed spinal cord is shown in figure 1(b).

**Figure 1.**
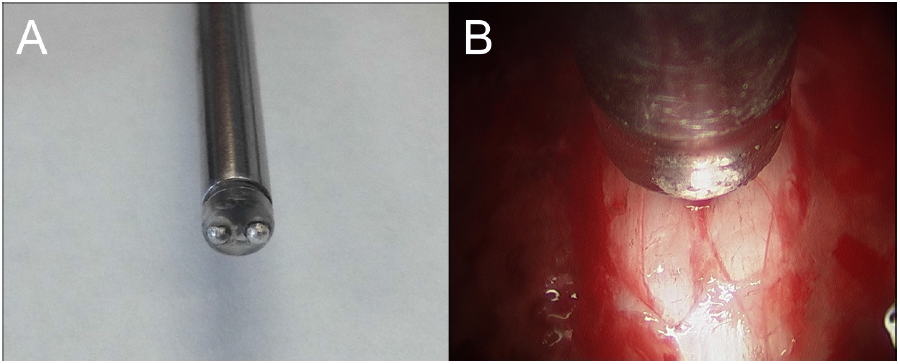
(a) Distal tip of the bipolar probe, showing the two hemispherical electrodes of 1 mm diameter each. (b) Intraoperative neuro-endoscopic photograph of the distal tip of the probe in place on the pial surface of the spinal cord.

### 2.2 Stimulation protocol

The essence of the experiments was to capture the waveform of an AC stimulation signal, *V*, applied across the electrodes, while simultaneously capturing the waveform of the resulting current, *I*, flowing through the electrode-tissue-electrode loop. By measuring current as a function of the frequency of the stimulation signal, we could then resolve the resistive and reactive components of the electrode-tissue impedance over the entire range of frequencies of interest for clinical spinal cord stimulation and beyond. For this purpose, we made our measurements in a 1-2-5 logarithmic sequence of stimulation frequencies over the nominal range from 1 Hz to 1 MHz, using both sine and square waves at several intensities between 50 mV and 5 V peak-to-peak. The study consisted of two arms: the *in vivo* stimulation experiments, and a series of *in vitro* system characterization experiments. The latter were done to establish the baseline performance of the measurement system, to evaluate the effects of noise in the data, and to confirm front-to-back calibration and linearity of the instrumentation chain. A summary of the measurements made in both arms of the study is shown in table 1. The *in vivo* experiments were carried out in the translational research facilities of the U.I. Department of Orthopedics and Rehabilitation, and the bench top *in vitro* work was done in the laboratories of the U.I. Department of Neurosurgery. The study design and data analysis tasks took place primarily at the Universities of Southampton and Virginia.

**Table 1.** Overview of the protocol for the *in vivo* ovine studies and the supplementary bench top *in vitro* and noise measurement tests.

| ***In Vivo Tests*** |  |  |  |
| --- | --- | --- | --- |
| *Sheep Identifier Number* | *Number of Data Blocks* | *Waveforms Used* | *Electrode Pair Orientations* |
| 1 | 8 | 7 sine wave<br>1 square wave | 8 runs parallel |
| 2 | 10 | 8 square wave<br>2 sine wave | 5 runs parallel<br>5 runs perpendicular |
| 3 | 8 | 8 sine wave | 3 runs parallel<br>5 runs perpendicular |
| ***In Vitro Tests*** |  |  |  |
| Preliminary gel tests | 1 | 1 sine wave | N/A |
| Shunt resistor gel calibrations | 4 | 4 sine wave | N/A |
| Oscilloscope noise tests | 47 | sine waves and square waves | N/A |

### 2.3 Instrumentation and measurement procedures

A schematic diagram of the experimental arrangement is shown in figure 2. The source of the stimulation signals for the clinical work and for the *in vitro* gel studies was a precision Agilent model 33500B waveform generator. The initial design studies and the preliminary bench top tests with an artificial load were done with a BK Precision 4012A function generator. The measurements were made with a Keysight Technologies DSO1022A digital storage oscilloscope (200 MHz bandwidth) using a pair of Keysight Technologies N2791A matched, offset-nulled and battery-isolated 10:1 differential probes (25 MHz bandwidth).

**Figure 2.**
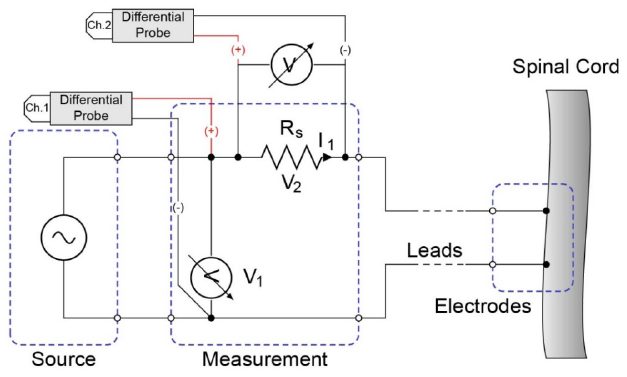
Block diagram showing the how the differential probes were used to measure the voltage across the electrodes on the spinal cord, and that across the shunt resistor thus determining the current flowing through the electrode-tissue-electrode loop.

Per figure 2, the voltage drop across the electrodes on the spinal cord was monitored on Channel 1 of the oscilloscope, and the voltage drop across a precision (1%) shunt resistor *R_S_* in series with the electrodes was monitored on Channel 2. The latter thus provided a measurement of the current flowing through the electrode-tissue-electrode loop. To help evaluate system response over the course of the *in vivo* experiments, shunt resistors with values of 100 Ω, 300 Ω, 500 Ω, 700 Ω and 1 kΩ were used. The oscilloscope’s signal averaging algorithm enabled noise reduction over most of the frequency range of interest, and was typically employed for stimulations from 50 Hz through 1 MHz. The sampling rates were adjusted as a function of the stimulation signal’s frequency in order to maximize the number of waveforms recorded during each stimulus presentation *vs.* the storage depth of the oscilloscope (10^4^ points per channel). All of the experimental data that constituted the waveform measurements were exported from the oscilloscope as *.CSV files for subsequent analysis. These were accompanied by *.TXT files documenting the relevant oscilloscope settings in each case.

### 2.4 System performance tests, noise measurements and in vitro studies

Assessment and confirmation of the measurement system’s technical performance was obtained using artificial loads to simulate the *in vivo* impedance. The two primary tests of this type were (1) measurement of the voltage drop across the shunt resistor as a function of frequency for stimulation at constant signal strength and with fixed load parameters, and (2) a comprehensive evaluation of the noise in the measurement system. The first type of test allowed us to anticipate the presence of resistive and reactive components of impedance that would be expected in the *in vivo* measurements. The nominal values of the components for this first test were *RS* = 100 Ω, *RL* = 26.4 kΩ, and *CL* = 10 nF, with sinusoidal stimulation at 3.5 Vrms. The values for the load components were rough *a priori* estimates of the maximum *in vivo* parameters. In the second type of test, the background noise level was measured at each point where a cable, lead or device was added to the circuit, with and without application of a stimulation signal. Representative results of these tests are presented in Section 3.

Prior to starting the *in vivo* experiments, the full measurement system including the bipolar electrode probe was tested using saline gel as a surrogate for spinal cord tissue. For preparation, 0.45 g NaCl was mixed into 495.5 ml of distilled water to make 0.9% normal saline solution, which was then boiled with 8 g of commercial-grade gelatin powder and subsequently refrigerated until the gel set. This phantom material resembled the gel-based [33] and polymeric [34] spinal cord surrogates we have developed in the past for other purposes, but with the saline allowing it to approximate physiological conductivities. The distal tip of the bipolar electrode probe was then lowered onto the surface of the gel and measurements were made using sinusoidal stimulation signals at amplitudes of 1, 2 and 5 V across the previously mentioned range of frequencies, with *RS* = 100 Ω. The results, presented in Section 3 below, verified the overall experimental approach, and provided a calibration for the measurements, in that the saline used for the phantom had well-established ionic properties and yielded stable, repeatable values of impedance.

### 2.5 In vivo Data processing

The *.CSV files exported from the oscilloscope were archived and then the data were imported into Mathematica (Version 10, Wolfram Inc.) for processing and analysis using custom-built routines. The current flowing through the electrode-tissue-electrode loop was computed from the measured voltage drop across the nominal value of the shunt resistor used in a particular data series. The exact frequency for each sinusoidal run was determined by computing a discrete Fourier transform of the measured trace and identifying the position of the first major peak. This yielded frequencies within less than 1% of the nominal value to which the waveform generator was set in all cases. The phase of the same peak was used to compute the phase difference, φ, between the current and voltage signals. The amplitude ratios, and hence the impedances, were determined from the respective peak heights.

## 3. Results

### 3.1 Technical performance tests

The results from the technical performance evaluations of the measurement system using a simple parallel RC network as an artificial load are shown in figure 3 and confirm satisfactory operation of the system. The shunt resistor voltage increases as a function of stimulation frequency reflecting the high-pass filtering nature of the circuit. The estimated shunt voltage as derived from the elementary circuit model (mentioned above) using the nominal values of the components is also shown in figure 3 and closely mirrors the experimental performance, as expected.

**Figure 3.**
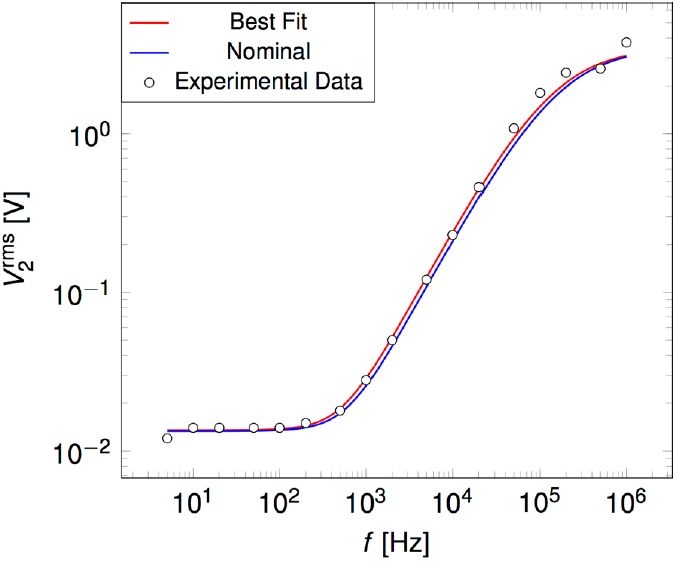
The measured shunt voltage vs. frequency for an RC network artificial load as compared with an elementary circuit theory prediction using the nominal values of the components. The expected high-pass filtering behavior was confirmed.

The noise measurements described in Section 2.4 were carried out in order to establish the baseline performance of the measurement system. Typical values of the signal-to-noise ratio (SNR) for the un-averaged voltage across the electrodes and for the current through the electrode-gel-electrode arrangement were 15.5 and 5.8, respectively. Figure 4 shows examples of the sweeps for the stimulation signal (blue trace) and the current response (green trace), with expanded views of the high-frequency noise components shown as inserts. Signal averaging during data acquisition increased the SNR values substantially and, as mentioned above, subsequent employment of the FFT analysis algorithm essentially eliminated any residual uncertainties in assessment of frequencies.

**Figure 4.**
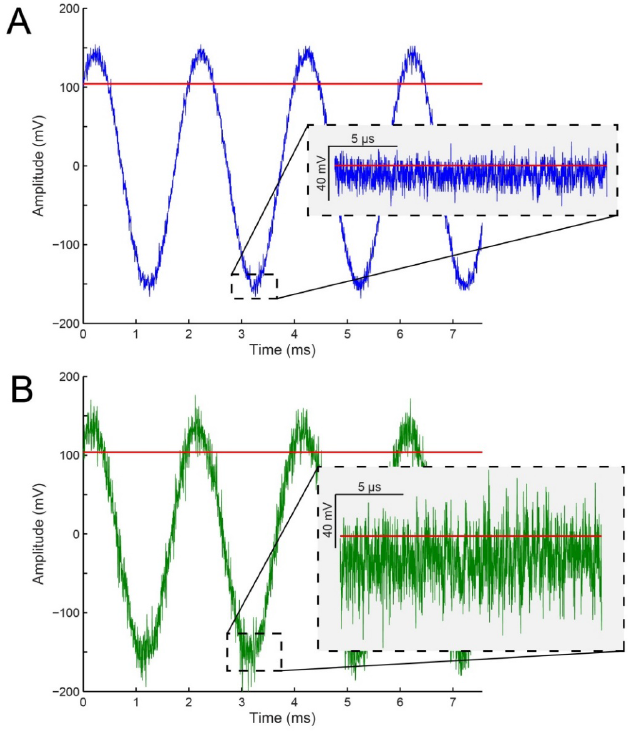
The un-averaged 500 Hz stimulation signal (A) and current response (B) as measured across a 1 kΩ shunt resistance during a full-system noise evaluation using a saline gel surrogate for the spinal cord tissues. Each plot’s inset depicts a brief portion of the waveform with its mean trend removed. The red lines indicate the displayed waveform’s RMS value.

### 3.2 In vitro studies

The results of the impedance vs. stimulation frequency measurements using the saline gelatin as the spinal cord tissue surrogate are shown in figure 5. The magnitude of the complex impedance trends downward over the ≈ 1 Hz to 1 MHz measurement range for all three of the stimulation amplitudes tested, with the phase decreasing monotonically at frequencies above 1 kHz. The impedances are higher at lower stimulation intensities, but converge to a common value of ≈ 200 Ω at stimulation frequencies near 1 MHz.

**Figure 5.**
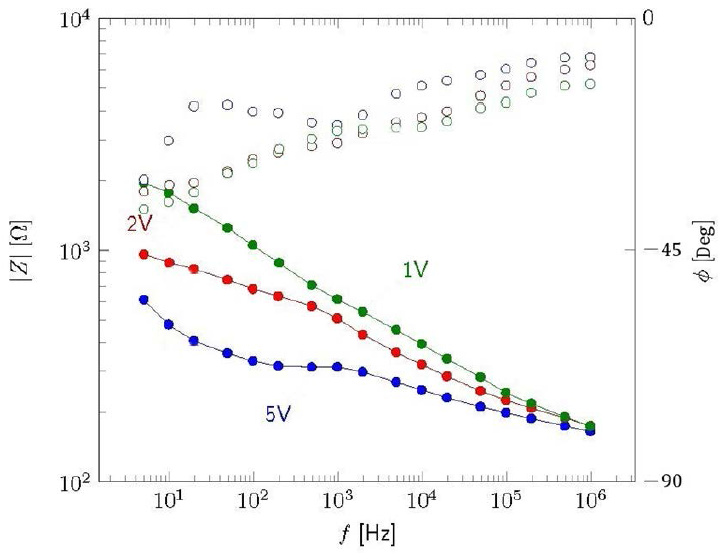
Complex impedance (solid circles) and phase (open circles) as a function of stimulation frequency as measured in a calibration study using the saline gelatin spinal cord surrogate, at three different stimulation signal strengths.

These findings approximate the corresponding features of the *in vivo* results presented below, thus demonstrating the utility of fixed-concentration saline gel as a robust and readily available phantom for system calibrations and preliminary tests. Moreover, because the gelatin is a homogeneous and isotropic material, the stimulation results are independent of the placement and alignment of the electrode pair on its surface. It thus serves as a useful baseline for confirming any directionality effects in spinal cord stimulations that might arise due to the axial orientation of the nerve fibers.

### 3.3 In vivo studies

Pilot trials of the experimental arrangement were carried out on Sheep No. 1 using only the parallel-electrode configuration. Full studies employing both the parallel- and perpendicular-electrode configurations were then carried out on Sheep Nos. 2 and 3. A total of 26 *in vivo* experiments were done across all three sheep. Examples of the raw data from one of them (Sheep 1, Block 6) are shown in figure 6, which includes plots of both un-averaged (20 Hz) and signal-averaged (2 kHz and 200 kHz) wave trains. As seen in Fig. 6, the presence of the frequency-dependent phase shift, φ, between the stimulation signal, V, and the resulting current, I, (represented by the response voltage across the shunt resistor) is evident from visual inspection.

**Figure 6.**
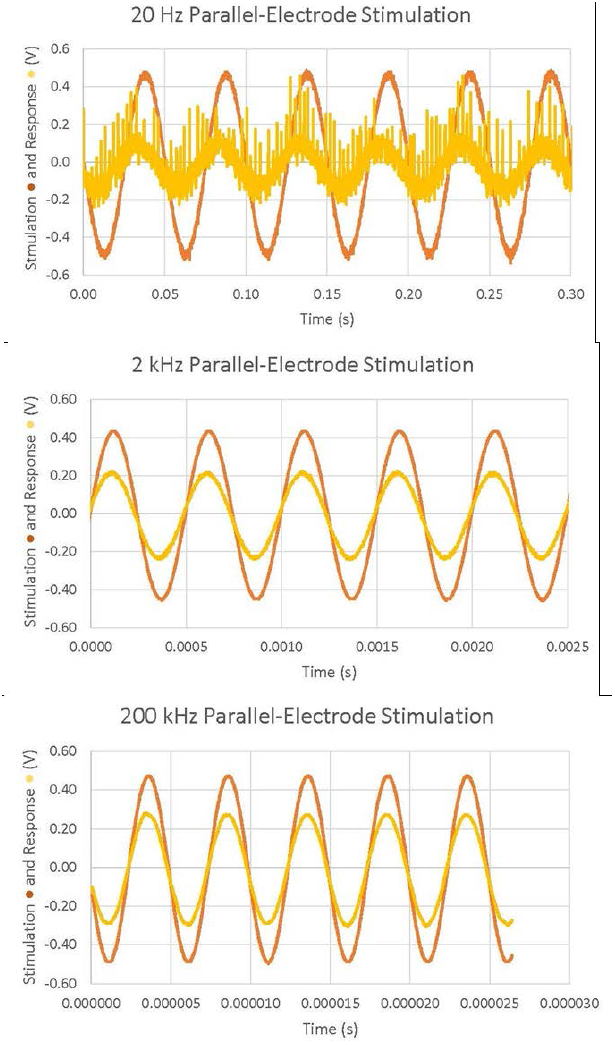
Examples of the raw-data wave trains acquired at 20 Hz, 2 kHz and 200 kHz for the case of *in vivo* sinusoidal stimulation at an amplitude of 500 mV with RS = 500 Ω and the electrodes in the parallel configuration. The 20 Hz data had no signal averaging; the data at 2 kHz and 200 kHz had 64x signal averaging.

Using the method described above, the magnitude and phase of the tissue impedance was obtained as a function of frequency. The results are shown in figures 7 and 8 respectively for all of the sinusoidal stimulation experiments that made direct comparisons of the parallel- and perpendicular-electrode configurations. In particular, this involved 8 data blocks from Sheep No. 3 with the resulting error bars representing a 95% confidence interval, and 1 data block from Sheep No. 2 (no error bars). As expected due to the fibrous structure of the spinal cord tissue, the impedance is larger in the case of the perpendicular arrangement, by about 50%. For instance, at a measured frequency of 200 Hz, the impedance is about 800 Ω in the parallel and 1300 Ω in the perpendicular direction. At these low frequencies, the impedance contains a strong negative reactance, as shown by the negative phase (figure 8). Interestingly, this is even a little more pronounced in the parallel than in the perpendicular arrangement.

**Figure 7.**
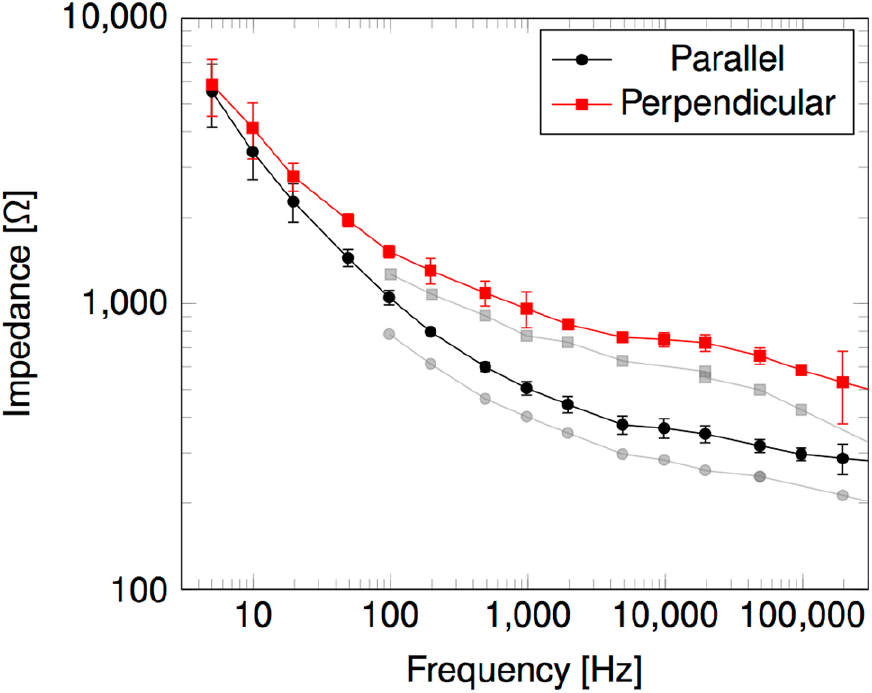
Magnitude of the complex tissue impedance vs. stimulation frequency for two sheep in both the parallel (solid circles) and perpendicular (solid squares) configurations of the pial surface electrodes. Sheep 3 (●,■), Sheep 2 (●,■).

**Figure 8.**
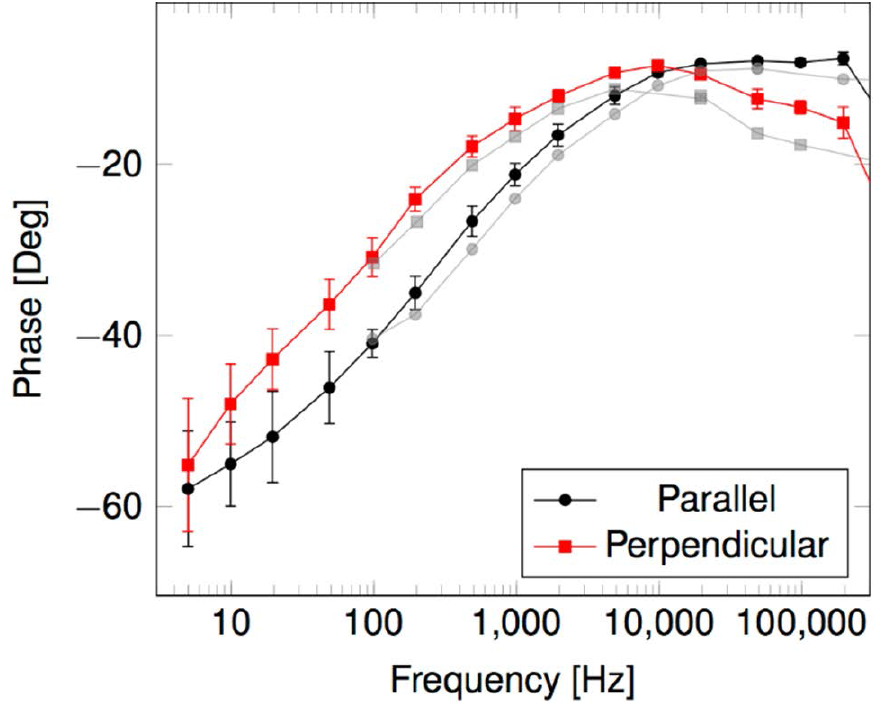
Phase of the complex tissue impedance vs. frequency for two sheep in both the parallel (solid circles) and perpendicular (solid squares) configurations of the pial surface electrodes. Sheep 3 (●,■), Sheep 2 (●,■).

Figures 9 and 10 show a particular example of these data in the form of the resistance and reactance (i.e., the real and imaginary parts of the complex impedance), respectively. The reactance, and with it the absolute value of the impedance, decreases steadily in magnitude as the frequency increases. However, there is a marked difference in behavior between the two orientations at high frequencies: while in the parallel direction, the reactance drops to a very small magnitude (about —10 Ω), it saturates at a significantly larger magnitude (—150 Ω) in the perpendicular case. This means that the tissue characteristics retain more of a capacitive nature at high frequencies if the electric field is applied perpendicular to the spinal cord axis.

**Figure 9.**
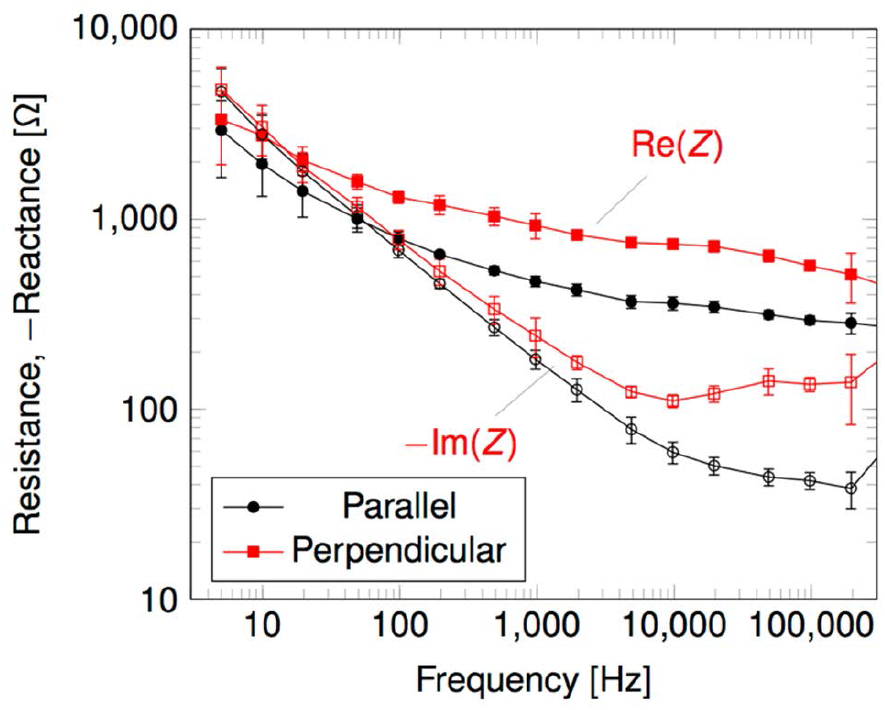
Representative example of the resistance (real, filled symbols) and reactance (imaginary, open symbols) component of the impedance for both the parallel and perpendicular configurations of the pial surface electrodes.

**Figure 10.**
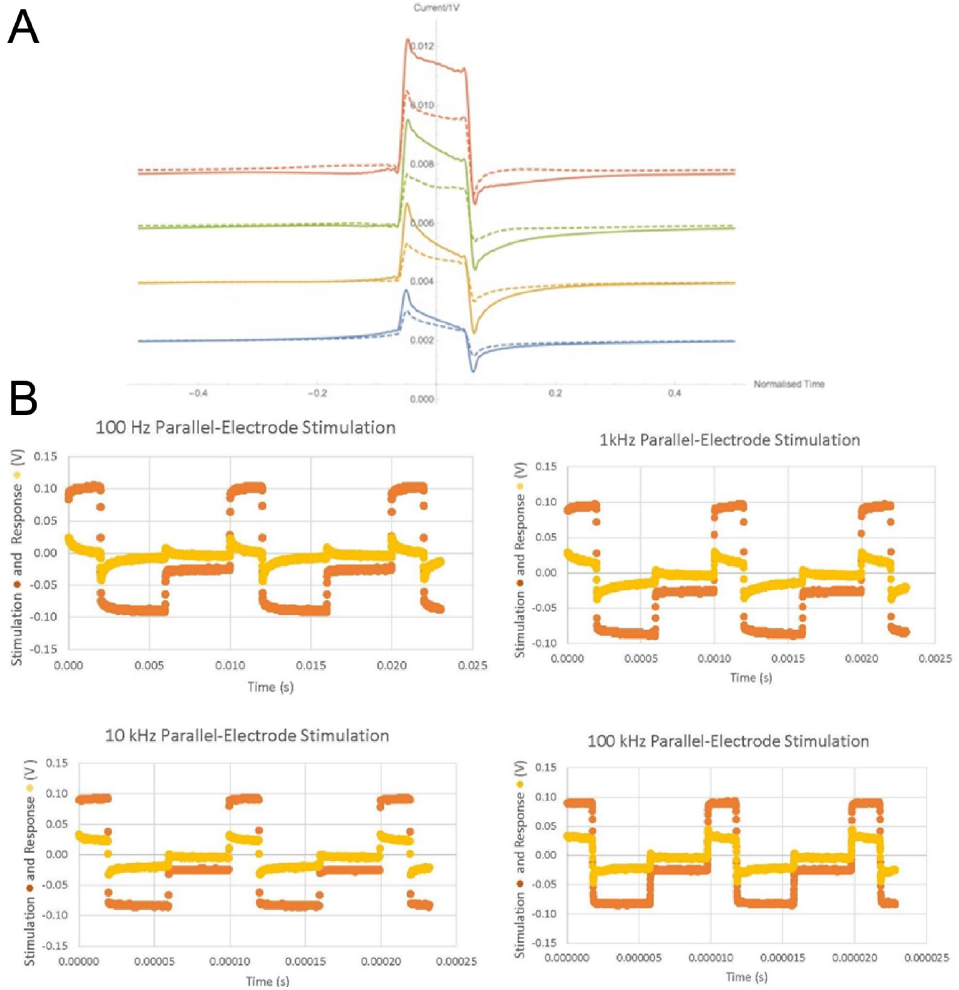
(a) Predicted current waveform response to a square unipolar voltage input of 10% duty ratio. The frequency increases from 10 Hz (bottom trace) to 100 kHz (top trace) by factors of 10. (b) Measured 0.1 Vpeak stimulation signals (●) and current response signals (●) for square wave excitation at 100 Hz, 1 kHz, 10 kHz and 100 kHz, showing increased fidelity of the response signal to the stimulation signal at higher frequencies.

## 4. Discussion

### 4.1 Optimization of stimulation strategies

#### 4.1.1 Intradural vs. epidural power delivery

The implications of these findings can be discussed quantitatively in terms of the power delivered to the target tissues under different SCS scenarios. The average power, *P*, delivered into the spinal cord by the electrodes is given by:

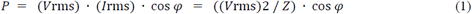

where *V*_rms_ and *I*_rms_ are the root-mean-square values of *V* and *I*, the impedance is *Z*, and cos *φ* is the power factor. From figure 8, *φ* decreases by roughly 15º per decade of frequency from 5 Hz to the plateau levels for both the parallel and perpendicular alignment cases, and there is ≈ 10º of offset between them until the phases equalize at about 10 kHz. At that point, cos *φ* achieves its maximum value of ≈ 0.99. From figure 7, for Sheep 3 and a mid-range stimulation amplitude of 0.5 V, the impedances for the parallel and perpendicular alignments of the electrodes are ≈ 400 Ω and 800 Ω at 10 kHz, and hence the average powers are 310 μW and 155 μW, respectively. At lower stimulation frequencies, e.g., the more clinically standard 100 Hz, the power levels associated with the parallel and perpendicular electrode alignments decrease to 95 μW and 72 μW respectively at the same stimulation amplitudes because the reactive component of the impedance is much higher at that frequency (see figure 9). However, it is important to note that while clinical stimulation is often performed at 100 Hz, the waveform is very different from a sinusoidal signal. In fact, the very short pulses that are employed contain significant Fourier components in the vicinity of 1/pulse width, not only at 1/pulse spacing.

There are no known fully equivalent data for the case of epidural SCS, i.e., studies in which complex impedance and phase were measured as a function of frequency of an epidural stimulation signal, and the resulting intraparenchymal power levels derived. At present, that includes our own case, in part because there is virtually no subdural space between the dura mater and the pial surface of the spinal cord in sheep, thus potentially confounding epidural vs. intradural SCS results. (However, see Section 4.3 below.) Perhaps the closest related studies are those of Alò et al. [30] and Abejon et al. [31]. In the former, a custom-made ohmmeter system was used to measure a mean bipolar impedance of 547 Ω during stimulation with 1 mA pulses between adjacent contacts in an epidural array in patients. They framed their discussion of the experimental design in terms of the energy consumption of the implanted pulse generator, appealing to the instantaneous power dissipated by the epidural load, which in their case would thus have been 547 μW. The latter authors measured the parameters of the perception threshold in SCS patients implanted with epidural leads, and found a mean impedance of 548 Ω associated with a mean stimulation current of 4.3 mA. These authors also employed ohmic modeling, in their case to determine therapeutic range of the stimulation process. From their reported parameters, the instantaneous epidural power dissipation would have been ≈ 10 mW.

To the extent that our complex-impedance intradural findings and the available ohmic-impedance epidural findings can be compared, direct intradural stimulation as tested here can operate at high frequencies in an almost fully resistive mode at power levels well below those reported for standard epidural stimulation. This suggests the possibility of extended battery life in clinical devices using the intradural approach.

#### 4.1.2 Electrode orientation

With direct high frequency stimulation, the orientation of the electrodes relative to the rostral-caudal axis of the spinal cord has a major influence on the impedance. By employing different bipolar montages of an intradural stimulation array, it should be possible to exploit these position-dependent impedance differences in order to produce volumetrically selective patterns of activation that are beyond the capabilities of epidural stimulation at any frequency. We anticipate that this will translate into a substantial difference in the functional and clinical capabilities of direct stimulation as compared with the standard methods. This difference arises because in epidural stimulation there is a relatively large distance separating the stimulus-producing electrodes and the neural structures targeted in the spinal cord. That space typically includes epidural fat, the dura mater and, most importantly, a thick layer (3 to 6 mm) [35] of CSF that has a large isotropic ionic conductivity which dominates the impedance. The CSF diffuses the epidurally-delivered stimulus patterns [1] thus limiting the spatial selectivity of activation. While substantial scientific, clinical and technical development programs have led to many improved epidural device designs and stimulation strategies [36,37,38], it is not clear that this fundamental physiological barrier to target selectivity can be definitively circumvented except through intradural placement of the electrode array. This situation is reflected in even the favorable findings of recent high frequency (10 kHz) epidural stimulation trials [16,17,18], in that the outcomes did not depend strongly on precise placement of the leads, which were implanted so as to generally cover the T8-T11 vertebral levels at the approximate midline.

#### 4.1.3 Waveform fidelity

The frequency dependence of the impedance as observed here also has consequences in how the tissue responds to rectangular voltage pulses, which are typically used in spinal cord stimulation. This is illustrated in figure 10. The current response of a unipolar-voltage pulse has been calculated and displayed in figure 10(a) at different frequencies based on the observed impedance. In all cases, the square wave input is distorted by the reactive component of the tissue impedance. However, the distortion is much more pronounced at low frequencies. The strong reactive nature of the tissue impedance essentially acts as a differentiator at low frequency, and the square unipolar voltage pulse results in a bipolar pair of current pulses that coincide with the rising and falling flanks of the voltage. At high frequencies, this transient behavior is still somewhat present, but the response is dominated by a unipolar square-current pulse. This behavior was observed in our *in vivo* experiments in which square-wave excitation was used. An example is shown in figure 10(b), where differentiation and attenuation of the current response pulse at low frequencies is evident upon visual inspection of the waveforms.

This has important implications for SCS in that the most effective waveforms will likely be those that resemble axonal action potentials. Action potential voltage changes are typically brief (about 1 to 2 ms duration) and have sharp rise and fall times. Because of the frequency-dependent impedance properties of the spinal cord tissue, high frequency pulse trains consisting of square wave electrical stimuli meant to mimic some aspects of action potentials will have their waveforms preserved. This should result in more effective activation of targeted neural structures than would occur with distorted lower frequency stimuli.

Others are exploring different approaches to the optimization of stimulation waveforms. Some examples include the genetic algorithm-based method of Wongsarnpigoon and Grill [39], the least-action principle pulses of Krouchev *et al* [40], and the patient-specific waveform synthesis of Yalçinkaya and Erbaş [41]. An important goal of such work is the minimization of pulse energy (thus obtaining extended battery life) while maximizing physiological effectiveness [42].

### 4.2 Implications for modeling of spinal cord stimulation

In a previous study, we used finite element modeling (FEM) to compare the electrical current distributions and the spatial recruitment profiles produced by direct intradural and epidural stimulation [43]. The results from that work, as well as those obtained by Howell et al [44], showed that direct intradural stimulation achieved greater selectivity in activating targeted structures within the spinal cord, and that it could do so at significantly lower power levels. That work employed a three-dimensional volume conductor approximation for the spinal cord tissues and assumed DC excitation of them, per the standard approach as reviewed by Holsheimer [9]. The impedance measurements reported here will be helpful in extending models of this kind to the regime of AC stimulation, since they provide a quantitative basis when comparing to predictions of electrodynamic current distributions made as a function of excitation frequency, signal strength, and electrode orientation.

Several other groups are also working towards improved models of the spinal cord stimulation process. These include the low-frequency dorsal horn network study of Zhang et al [45], the hybrid-biophysical computational approach of Capograsso *et al* [46], the improved dendritic channel method of Elbasiouny and Mushahwar [47], and the frequency-dependent interpolation approach of Tracey and Williams [48]. Modeling of these kinds provide very useful insights into the design and behavior of the stimulation electrodes [49], lead configurations [50,51,52,53], the synthesis of robust stimulation control algorithms [54], improved basic knowledge of nerve physiology [55], and possible mechanisms of action at work in high frequency epidural stimulation [56].

### 4.3 Ongoing efforts and future work

There is at present no spinal cord equivalent of a direct brain stimulation (DBS) system, in that virtually all of the SCS devices now in clinical use are implanted in the epidural space and thus encounter fundamental performance limits dictated by the presence of the intervening layer of CSF. The approach that we have proposed and are investigating [57] seeks to circumvent this limitation through direct intradural stimulation, and to re-introduce that method [58] as a clinical option employed against intractable neuropathic pain and the spasticity arising from spinal cord injury [59]. This would represent a significant departure from the *status quo*, wherein intradural placement of a stimulator lead is either an inadvertent result of attempted epidural placement [60] or, rarely, an intentional means of having the body of the lead serve as a dural substitute [61]. Our *in vivo* measurements of the frequency-dependent impedance of the spinal cord tissues provide the start of a quantitative basis for exploring direct intradural stimulation strategies, in much same way that similar studies [62] have opened windows onto the performance of DBS systems.

We are pursuing a number of improvements in our approach to these measurements aimed at overcoming existing limitations in them and extending the utility of the findings. These include possible measurement of the epidural impedance when the electrodes are placed on the surface of the dura mater. Data of that kind would enable quantitative assessment of how the presence of the CSF layer diffuses the stimulation signals, with the expected outcome being essentially no difference in the impedance patterns for parallel and perpendicular placement of the electrodes. However, as noted above, the subdural space in the sheep is almost non-existent. Hence an epidural placement of the electrodes on the dura mater without having the distal tips of their hemispherical surfaces causing it to make contact with the pia mater would be challenging. Even so, we are investigating ways to accomplish a purely epidural placement of the electrodes via careful micromanipulation of the probe. The limiting factor in achieving a stable separation between the dura and pia in that situation will likely be the intrinsic diametric pulsations (≈ 100 μm in amplitude) of the spinal cord associated with the cardiac cycle [63]. An alternative approach to a comparative intradural/epidural measurement would simply be to create a window in the dura, perform direct stimulation, withdraw the electrode probe, fill the cavity with artificial CSF and then stimulate.

We are also developing neurophysiological methods for obtaining single unit recordings from dorsal horn neurons identified as being those most responsive to spinal cord stimulation signals [64]. The protocols for experiments of that kind now in progress will be expanded to include simultaneous measurement of spinal cord impedance while the single unit recordings are being obtained from targets at different depths within the dorsal horn, and with different stimulus presentations (eg., frequency sweeps as described in Section 3, burst patterns, different waveforms, etc.). Results from a study of this kind would provide very useful validation data for FEM-based attempts at optimizing intradural stimulation patterns in order to maximize therapeutic efficacy.

Lastly, the ideal experimental arrangement would be one which enabled a chronic study in awake animals during their normal activities, in order to eliminate any potential systematic effects in the data that might arise from the anesthesia. However, that would require an implantable pulse generator system that would be able to record and wirelessly report the stimulation voltage and current signals.

## 5. Conclusions

In summary, we have measured the effective tissue impedance of the ovine spinal cord *in vivo* as it presents itself to a pair of stimulation electrodes placed directly in contact with the pial surface, in orientations both parallel and perpendicular to the rostral-caudal axis. Generally, the impedance is smaller for high frequency components, and becomes more resistive rather than capacitive in nature. This is to be expected due to the ionic conductance of the tissue, with ion mobility restricted by the tissue structure. As a consequence, low frequency (≈ 100 Hz) stimulation signals result in highly distorted current responses. At high frequencies (≈ 10 kHz and above), the tissue response is mostly resistive and the current waveforms maintain fidelity with those of the source stimulation signals.

## Acknowledgments

The authors thank their University of Iowa colleagues H Oya and B D Dalm for their interest in this work and for useful discussions. Co-authors Utz, Reddy, Gillies and Howard may receive patent royalties from any commercial licensing of the I-Patch intradural stimulator intellectual properties negotiated by the Universities of Iowa and Virginia.

## References

1. Holsheimer J 2002 Which neuronal elements are activated directly by spinal cord stimulation Neuromodulation 5 25–31.

2. Bradley K 2006 The technology: the anatomy of a spinal cord and nerve root stimulator: the lead and power source Pain Med. 7 S27–S34.

3. North RB 2008 Neural interface devices: spinal cord stimulation technology Proc. IEEE 96 1108–19.

4. Levy R M 2013 Progress in the technology of neuromodulation: the emperor’s new clothes? Neuromodulation 16 285–91.

5. Holsheimer J, Struijk J J and Rijkhoff N J M 1991 Contact combinations in epidural spinal cord stimulation Stereotact. Func. Neurosurg. 56 220–33.

6. He J, Barolat G, Holsheimer J and Struijk J J 1994 Perception threshold and electrode position for spinal cord stimulation Pain 59 55–63.

7. Holsheimer J and Wesselink W A 1997 Optimum electrode geometry for spinal cord stimulation: the narrow bipole and tripole Med. Biol. Eng. Comput. 35 493–7.

8. Barolat G 1998 Epidural spinal cord stimulation: anatomical and electrical properties of the intraspinal structures relevant to spinal cord stimulation and clinical correlations Neuromodulation 1 63–71.

9. Holsheimer J 1998 Computer modelling of spinal cord stimulation and its contribution to therapeutic efficacy Spinal Cord 36 531–40.

10. Holsheimer J, Khan Y N, Raza S S and Khan E A 2007 Effects of electrode positioning on perception threshold and paresthesia coverage in spinal cord stimulation Neuromodulation 10 34–41.

11. De Vos C C, Hilgerink M P, Buschman H P J, and Holsheimer J 2009 Electrode contact configuration and energy consumption in spinal cord stimulation Neurosurgery 65(S6) 210–6.

12. Ackermann JrD M, Bhadra N, Foldes E L, and Kilgore K L 2011 Conduction block of whole nerve without onset firing using combined high frequency and direct current Med. Biol. Eng. Comput. 49 241–51.

13. Cuellar J M, Alataris K, Walker A, Yeomans D C, and Antognini J F 2013 Effect of high-frequency alternating current on spinal afferent nociceptive transmission Neuromodulation 16 318–27.

14. Fisher K M, Jillani N E, Oluoch G O, and Baker S N 2015 Blocking central pathways in the primate motor system using high-frequency sinusoidal current J. Neurophysiol. 113 1670–80.

15. Reddy C G, Dalm B D, Flouty O E, Gillies G T, Howard IIIM A, and Brennan T J Comparison of conventional and kilohertz frequency epidural stimulatin in patients undergoing trialing for spinal cord stimulation: clinical considerations World Neurosurg. in press.

16. Van Buyten JP, Al-Kaisy A, Smet I, Palmisani S, and Smith T 2013 High-frequency spinal cord stimulation for the treatment of chronic back pain patients: results of a prospective multicenter european clinical study Neurmodulation 16 59–66.

17. Al-Kaisy A, Van Buyten J-P, Smet I, Palmisani S, Pang D, and Smith T 2014 Sustained effectiveness of 10 kHz high-frequency spinal cord stimulation for patients with chronic, low back pain: 24-month results of a prospective multicenter study Pain Med. 15 347–54.

18. Kapural L, Yu C, Doust M W, Gliner B E, Vallejo R, Sitzman B T, Amirdelfan K, Morgan D M, Brown L L, Yearwood T L, Bundschu R, Burton A W, Yang T, Benyamin R, and Burgher A H 2015 Novel 10-kHz high-frequency therapy (HF10 therapy) is superior to traditional low-frequency spinal cord stimulation for the treatment of chronic back and leg pain Anesthesiology 12 851–60.

19. Eldabe S, Kumar K, Buchser E andTaylor R S 2010 An analysis of the components of pain, function, and health-related quality of life in patients with failed back surgery syndrome treated with spinal cord stimulation or conventional medical Management Neuromodulation 13 201–9.

20. Kilgore K L and Bhadra N 2014 Reversible nerve conduction block using kilohertz frequency alternating current Neuromodulation 17 242–55.

21. Levy R M 2014 Anatomic considerations for spinal cord stimulation Neuromodulation 17(Suppl.) 2–11.

22. Howard IIIM A, Utz M, Brennan T J, Dalm B D, Viljoen S, Jeffery N D, and Gillies G T 2011 Intradural approach to selective stimulation in the spinal cord for treatment of intractable pain: design principles and wireless protocol J. Appl. Phys. 110 044702.

23. Foutz T J and McIntyre C C 2010 Evaluation of novel stimulus waveforms for deep brain stimulation J. Neural Eng. 7 06608.

24. Dalm B D, Viljoen S V, Dahdaleh N S, Reddy C G, Brennan T J, Oya H, Wilson S, Safayi, S, Jeffery N D, Gillies G T, and Howard IIIM A 2014 Revisiting intradural spinal cord stimulation: an introduction to a novel intradural spinal cord stimulation device Innov. Neurosurg. 2 13–20.

25. Oliynyk M S, Gillies G T, Oya H, Wilson S, Reddy C G, and Howard IIIM A 2013 Dynamic loading characteristics of an intradural spinal cord stimulator J. Appl. Phys. 113 026103.

26. Grosland N M, Gillies G T, Shurig R, Stoner K, Viljoen S, Dalm B D, Oya H., Fredericks D C, Gibson-Corley K, Reddy C, Wilson S, and Howard IIIM A 2014 Finite-element study of the performance characteristics of an intradural spinal cord stimulator ASME J. Medical Devices 8 041012.

27. Flouty O E, Oya H, Kawasaki H, Reddy C G, Fredericks D C, Gibson-Corley K N, Jeffery N D, Gillies G T, and Howard IIIM A 2013 Intracranial somatosensory responses with direct spinal cord stimulation in anesthetized sheep PLOS ONE 8 e56266.

28. Oya H, Safayi S, Jeffery N D, Viljoen S, Reddy C G, Dalm B D, Kanwal J K, Gillies G T, and Howard IIIM A 2013 Soft-coupling suspension system for an intradural spinal cord stimulator: biophysical performance characteristics J. Appl. Phys. 114 164701.

29. Wei X F and Grill W M 2009 Impedance characteristics of deep brain stimulation electrodes in vitro and in vivo J. Neural Eng. 6 046008.

30. Alò K, Varga C, Krames E, Prager J, Holsheimer J, Manola L, and Bradley K 2006 Factors affecting impedance of percutaneous leads in spinal cord stimulation Neuromodulation 9 128–35.

31. Abejon D and Feler C A 2007 Is impedance a parameter to be taken into account in spinal cord stimulation? Pain Physician 10 533–40.

32. Gibson-Corley K N, Oya H, Flouty O, Fredericks D C, Jeffery N D, Gillies G T, and Howard IIIM A 2012 Ovine tests of a novel spinal cord neuromodulator and dentate ligament fixation method Journal of Investigative Surgery 25 366–74.

33. Gillies GT, Wilhelm TD, Humphrey JAC, Fillmore HL, Holloway KL, and Broaddus WC 2002 Spinal cord surrogate with nanoscale porosity for in vitro simulations of restorative neurosurgical techniques Nanotechnology 13 587–591.

34. Howard IIIMA, Utz M, Brennan TJ, Dalm BD, Viljoen S, Kanwal JK, and Gillies GT 2011 Biophysical attributes of an in vitro spinal cord surrogate for use in developing an intradural neuromodulation system J. Appl. Phys. 110 074701.

35. Lee RA, van Zundert AAJ, Breedveld P, Wondergem JHM, Peek D, and Wieringa PA 2007 The anatomy of the thoracic spinal canal investigated with magnetic resonance imaging (MRI) Acta Anaesth. Belg. 58 163–7.

36. Oakley JC, Krames ES, Prager JP, Stamatos J, Foster AM, Weiner R, Rashbaum RR, and Henderson J 2007 A new spinal cord stimulation system effectively relieves chronic, intractable pain: a multicenter prospective clinical study Neuromodulation 10 262–78.

37. Molnar G and Barolat G 2014 Principles of cord activation during spinal cord stimulation Neuromodulation 17 12–21.

38. Kumar K, Caraway DL, Rizvi S, and Bishop S 2014 Current challenges in spinal cord stimulation Neuromodulation 17 22–35.

39. Wongsarnpigoon A and Grill WM 2010 Energy-efficient waveformshapes for neural stimulation revealed with genetic algorithm J. Neural Eng. 7 046009.

40. Krouchev NI, Danner SM, Vinet A, Rattay F, and Sawan M 2014 Energy-optimal electrical-stimulation pulses shaped by the lease-action principle PLoS One 9 e90480.

41. Yalçinkaya F and Erbaş A 2014 The design of an embedded spinal cord stimulator Turk. J. Elec. Eng. Comp. Sci. 22 1453–62.

42. Foutz TJ and McIntyre CC 2010 Evaluation of novel stimulus waveforms for deep brain stimulation J. Neural Eng. 7 066008.

43. Huang Q, Oya H, Flouty O E, Reddy C G, Howard IIIM A, Gillies G T, and Utz M 2014 Comparison of spinal cord stimulation profiles from intra- and extradural electrode arrangements by finite element modelling Med. Biol. Eng. Comput. 52 531–8.

44. Howell B, Lad S P, and Grill W M 2014 Evaluation of intradural stimulation efficiency and selectivity in a computational model of spinal cord stimulation PLoS ONE 9 e114938.

45. Zhang T C, Janik J J, and Grill W M 2014 Modeling effects of spinal cord stimulation on wide-dynamic range dorsal horn neurons: influence of stimulation frequency and GABAergic inhibition J. Neurophysiol. 112 552–67.

46. Capogrosso M, Wenger N, Raspopovic S, Musienko P, Beauparlant J, Bassi Luciani L, Courtine G and Micera S 2013 A computational model for epidural electrical stimulation of spinal sensorimotor circuits J. Neurosci. 33 19326–40.

47. Elbasiouny S M and Mushahwar V K 2007 Modulation of motoneuronal firing behavior after spinal cord injury using intraspinal microstimulation current pulses: a modeling study J. Appl. Physiol. 103 276–86.

48. Tracey B and Williams M 2011 Computationally efficient bioelectric field modeling and effects of frequency-dependent tissue capacitance J. Neural Eng. 8 036017.

49. McIntyre C C and Grill W M 2001 Finite element analysis of the current-density and electric field generated by metal microelectrodes Ann. Biomed. Eng. 29 227–35.

50. Manola J, Holsheimer J 2004 Technical performance of percutaneous and laminectomy leads analyzed by modeling Neuromodulation 7 231–41.

51. Sankarasubramanian V, Buitenweg J R, Holsheimer J, and Veltink P 2011 Electrode alignment of transverse tripoles using a percutaneous triple-lead approach in spinal cord stimulation J. Neural Eng. 8 016010.

52. Sankarasubramanian V, Buitenweg J R, Holsheimer J, and Veltink P 2011 Triple leads programmed to perform as longitudinal guarded cathodes in spinal cord stimulation: a modeling study Neuromodulation 14 401–11.

53. Levy RM 2014 Anatomic considerations for spinal cord stimulation Neuromodulation 17 2–11.

54. Denison T and Litt B 2014 Advancing neuromodulation through control systems: a general framework and case study in posture-responsive stimulation Neuromodulation, 17 48–57.

55. Cranford J P, Kim B J, and Krassowska Neu W 2012 Asymptotic model of electrical stimulation of nerve fibers Med. Biol. Eng. Comput. 50 243–51.

56. Lempka S F, McIntyre C C, Kilgore K L and Machado A G Computational analysis of kilohertz frequency spinal cord stimulation for chronic pain management Anesthesiology 122 1362–76.

57. Flouty O E, Oya H, Kawasaki H, Wilson S, Reddy C G, Jeffery N D, Brennan T J, Gibson-Corley K N, Utz M, Gillies G T, and Howard IIIM A 2012 A new device concept for directly modulating spinal cord pathways: initial in vivo experimental results Physiol. Meas. 33 2003–15.

58. Gibson-Corley KN, Flouty O, Oya H, Gillies GT, and Howard IIIMA 2014 Postsurgical pathologies associated with intradural electrical stimulation in the central nervous system: design implications for a new clinical device BioMed Res. Int. 2014 989175.

59. Safayi S, Jeffery ND, Shivapour SK, Zamanighome M, Zylstra TJ, Bratsch-Prince J, Wilson S, Reddy CG, Fredericks DC, Gillies GT, and Howard IIIMA 2015 Kinematic analysis of the gait of adult sheep during treadmill locomotion: parameter values, allowable total error, and potential for use in evaluating spinal cord injury J. Neurol. Sci. 358 107–112.

60. Pope JE and Stanton-Hicks M 2010 Accidental subdural spinal cord stimulator lead placement and stimulation Neuromodulation 14 30–3.

61. Menovsky T, De Ridder D and De Mulder G 2009 Placement of an electrode array as a dural substitute for dorsal column stimulation: technical note Minim. Invas. Neurosurg. 52 53–5.

62. Butson C R and McIntyre C C 2005 Tissue and electrode capacitance reduce neural activation volumes during deep brain stimulation Clin. Neurophys. 116 2490–2500.

63. Matsuzaki H, Wakabayashi K, Ishihara K, Ishikawa H, Kawabata H, and Onomura T 1995 The origin and significance of spinal cord pulsation Spinal Cord 34 422–6.

64. Miller JW, Reddy CG, Wilson S, Dalm BD, Safayi S, Shivapour SK, Abode-Iyamah K, Viljoen S, Fredericks DC, Gibson-Corley KN, Grosland NM, Jerat NU, Stoner K, Reale R, Oya H, Jefferey ND, Brennan TJ, Gillies GT, and Howard IIIMA 2015 Spinal cord stimulation in an ovine model of neuropathic pain measured through Von Frey filaments, gait analysis, and dorsal horn recording Abstract Book of the 3rd Annual Minnesota Neuromodulation Symposium (University of Minnesota, Minneapolis, Minnesota), p. 55 http://neuromodulation.umn.edu/2015NMProgram.pdf.

